# Serum cytokine levels are modulated by specific frequencies, amplitudes, and pulse widths of vagus nerve stimulation

**DOI:** 10.1101/2020.01.08.898890

**Authors:** Téa Tsaava, Timir Datta-Chaudhuri, Meghan E. Addorisio, Emily Battinelli Masi, Harold A. Silverman, Justin E. Newman, Gavin H. Imperato, Chad Bouton, Kevin J. Tracey, Sangeeta S. Chavan, Eric H. Chang

## Abstract

Electrical stimulation of peripheral nerves is a widely used technique to treat a variety of conditions including chronic pain, motor impairment, headaches, and epilepsy. Nerve stimulation to achieve efficacious symptomatic relief depends on the proper selection of electrical stimulation parameters to recruit the appropriate fibers within a nerve. Recently, electrical stimulation of the vagus nerve has shown promise for controlling inflammation and clinical trials have demonstrated efficacy for the treatment of inflammatory disorders. This application of vagus nerve stimulation activates the inflammatory reflex, reducing levels of inflammatory cytokines during inflammation. Here, we wanted to test whether altering the parameters of electrical vagus nerve stimulation would change circulating cytokine levels of normal healthy animals in the absence of increased inflammation. To examine this, we systematically tested a set of electrical stimulation parameters and measured serum cytokine levels in healthy mice. Surprisingly, we found that specific combinations of pulse width, pulse amplitude, and frequency produced significant increases of the pro-inflammatory cytokine tumor necrosis factor alpha (TNFα), while other parameters selectively lowered serum TNFα levels, as compared to sham-stimulated mice. In addition, serum levels of the anti-inflammatory cytokine interleukin-10 (IL-10) were significantly increased by select parameters of electrical stimulation but remained unchanged with others. These results indicate that electrical stimulation parameter selection is critically important for the modulation of cytokines via the cervical vagus nerve and that specific cytokines can be increased by electrical stimulation in the absence of inflammation. As the next generation of bioelectronic therapies and devices are developed to capitalize on the neural regulation of inflammation, the selection of nerve stimulation parameters will be a critically important variable for achieving cytokine-specific changes.

## INTRODUCTION

Electrical stimulation is a fundamental neuromodulation technique used to activate nerves and muscles within the body. When electrical current is applied to nerve tissue using electrodes, there are many multiple variables that determine whether the desired neural fibers become activated. Generally, fiber activation by electrical stimulation follows a recruitment order from largest to smallest, with the smallest diameter fibers requiring the highest stimulation current levels (Baratta et al., 1989; Blair et al., 1933; Fang and Mortimer, 1991). Because different fiber types in the peripheral nervous system have varying functions and innervate different target organs, this fiber recruitment principle is important to achieve the desired physiological changes from activating nerves (Gorman and Mortimer, 1983). Electrical stimulation parameters such as output frequency, current, duration, and amplitude, are important determinants for achieving effective nerve activation that is both selective and efficient (Grill, 2015).

In the vagus nerve, there are three main fiber types: A-, B-, and C-fibers that can be distinguished on the basis of axon diameters, myelination, conduction velocity and stimulation thresholds for activation (Heck et al., 2002; Groves and Brown, 2005). This mixed nerve carries sensory afferent and motor efferent signals between the brain and the body to mediate vital functions of the autonomic nervous system (Bonaz et al., 2013; Tracey, 2002). A large body of preclinical studies and emerging clinical evidence indicates that electrical stimulation at the cervical vagus nerve is able to change the body’s immune response to injury or infection, specifically by reducing the levels of certain serum cytokines that are important mediators of inflammation in the body (Andersson and Tracey, 2012; Borovikova et al., 2000). This stimulation of the vagus nerve regulates cytokine release from the spleen through activation of the *inflammatory reflex*, thereby protecting against lethality in models of systemic inflammation (Borovikova et al., 2000; Chavan et al., 2017; Tracey, 2002). Early clinical evidence for the therapeutic efficacy of this electrical vagus nerve stimulation has shown promise to treat patients with chronic inflammatory disorders (Bonaz et al., 2016; Koopman et al., 2016). This body of evidence has also demonstrated that electrical stimulation parameters can be modified to intentionally elicit different physiological effects, such as the separation of anti-inflammatory and cardioinhibitory effects (Huston et al., 2007). The anti-inflammatory effects of vagus nerve stimulation have been attributed to A- and B-fiber activation, while the cardioinhibitory effects are thought to be mediated by only B-fibers (Huston et al., 2007; Olofsson et al., 2015; Yoo et al., 2016).

As differential fiber recruitment is affected by nerve stimulation parameters, we wondered how specific stimulation parameters might effect serum cytokine levels in a parameter-dependent fashion. To accomplish this, we modified the output of our electrical stimulator to select frequencies, amplitudes, and pulse widths predicted to differentially activate different classes of fibers within the vagus nerve. Here we observed that certain cytokines, such as tumor necrosis factor alpha (TNFα) and interleukin 10 (IL-10), could be increased or decreased by specific combinations of frequency, pulse width, and pulse amplitudes in healthy mice.

## METHODS

### Animals

Naïve male BALB/c mice (8 to 12 weeks old) were obtained from Charles River Laboratories (Wilmington, MA, USA) and acclimated for at least one week before conducting experiments. Animals were housed on a 12:12 hour reverse light/dark cycle at 23° C and relative humidity 30-70%. Standard chow was withheld from animals for a period of up to three hours prior to stimulation of the cervical vagus nerve. Water was available *ad libitum*. All experiments were performed under protocols approved by the Institutional Animal Care and Use Committee of the Feinstein Institutes for Medical Research and in strict adherence with NIH guidelines on the care and use of laboratory animals.

### Vagus nerve isolation and electrical stimulation

All surgical procedures were conducted using aseptic technique. Mice were administered isoflurane anesthesia through a nose cone in the supine position (oxygen flow 1L/min, isoflurane 1.75%). An appropriate depth of anesthesia was assessed by toe pinch reflex. The cervical region was shaved with an electric razor and then sterilized with 70% ethanol. A midline incision was made, the salivary glands were identified and bluntly dissected to expose the left carotid bundle lateral to the sternocleidomastoid muscle. The left cervical vagus nerve was separated from the carotid sheath and placed on a 200 μm diameter micro cuff sling bipolar electrode with platinum-iridium contacts (CorTec GmbH, Freiburg, Germany). Parafilm was placed over the surgical site to prevent desiccation of the nerve during stimulation. Heart rate was continuously monitored using a MouseSTAT^®^ Heart Rate Monitor (Kent Scientific, Torrington, CT, USA).

Electrical pulses were delivered by a constant current stimulator system PlexStim 2.0 (Plexon, Dallas, TX) and individually controlled with PlexStim v2.2 software. The following stimulation parameters were used for stimulation: Four minute duration, pulse width (50μs, 250μs), amplitude (50μa, 200μA, and 750μA), and frequency (30Hz, 100Hz). Sham operated mice underwent the same surgical procedures but without electrical stimulation. After stimulation, the skin was sutured closed and the animals were returned to their home cages for recovery.

### Serum collection and analysis

Two hours following stimulation, whole blood was collected from animals by cardiac puncture following euthanasia by CO_2_ asphyxiation. Blood was allowed to clot in a polypropylene tube at room temperature for 30-60 min. To obtain serum, the tubes were centrifuged two times, first at 5000 x g for 10 minutes, followed by 10000 x g for 2 minutes. The supernatant serum was collected in a clean tube and stored at −20°C until further processing. Serum was analyzed on multiplex cytokine immunoassay plates (V-PLEX Panel 1 mouse kit; Meso Scale Discovery, Rockville MD) to quantify levels of IFN-γ, IL-1β, IL-2, IL-4, IL-5, IL-6, CXCL1, IL-10, IL-12p70, and TNF-α.

### Statistical analysis

Differences in serum cytokine levels between stimulated and non-stimulated (sham) groups were analyzed by Mann-Whitney U tests (Prism 8.0). In all tests, *P*<0.05 was accepted as an indication of statistical significance.

## RESULTS

Here we utilized an established methodology to surgically isolate and electrically stimulate the vagus nerve. For all experiments, stimulation was performed on experimental animals with sham-matched controls. The vagus nerve at the cervical level was surgically isolated and a cuff electrode was placed on the nerve. Following stimulation, the mouse was recovered for two hours, then euthanized for tissue collection (**Figure 1A**). During electrical stimulation, depolarization of the nerve was achieved via cathodal current application at the electrode-tissue interface and at the electrode with the lower electrical potential. We utilized biphasic waveforms, with a secondary anodic phase to balance out the net effect charge to zero (**Figure 1B**). Without charge balancing, a detrimental electrical potential buildup at the electrode may occur over time resulting in both tissue damage and reduction of electrode effectiveness. The stimulation waveforms for the different parameters are shown in **Figure 1B.**

**Figure 1.**
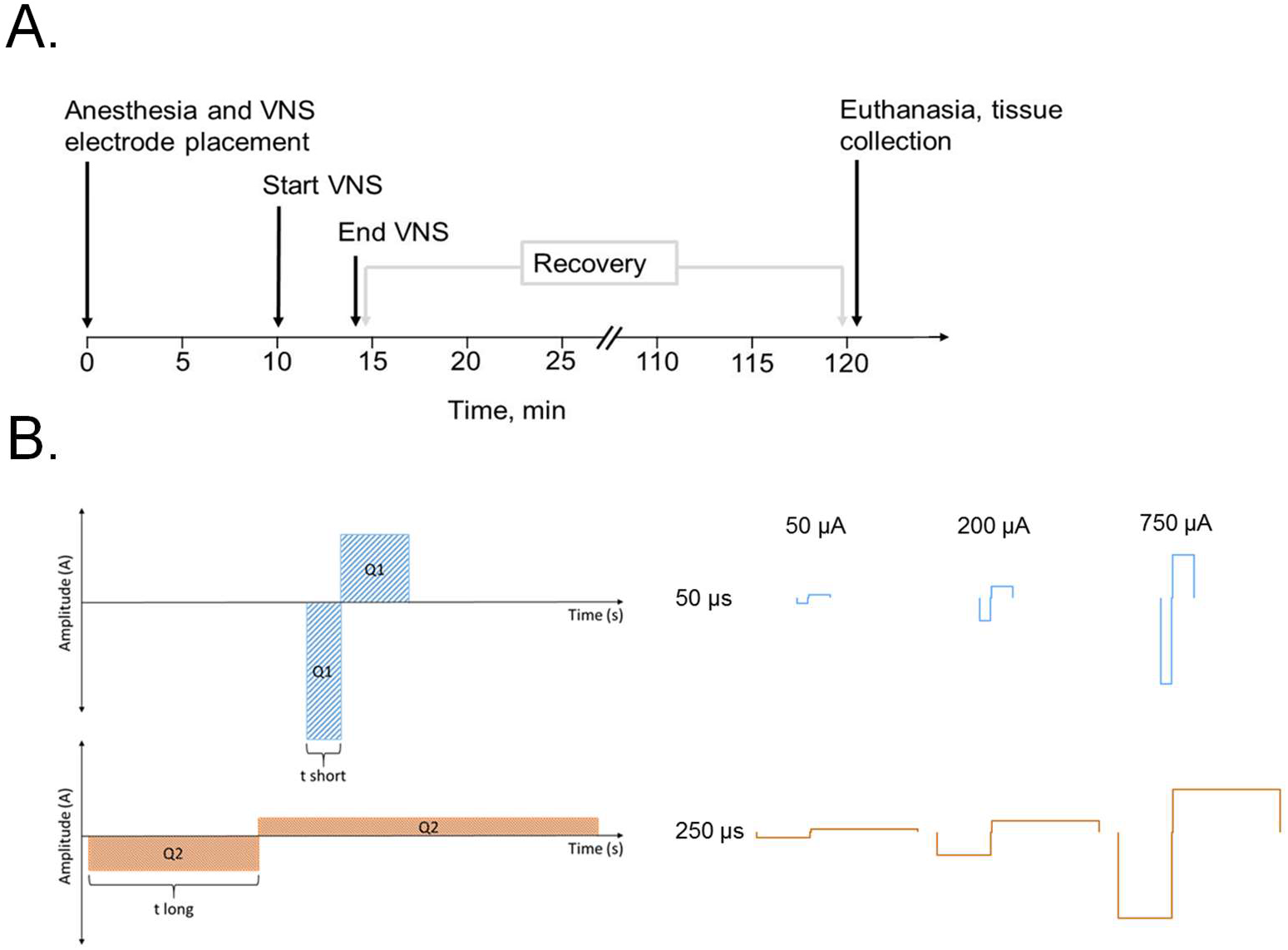
Experimental design and stimulation pulse waveforms. **A.)** Experimental timeline. Electrical stimulation pulse trains were applied to the exposed left cervical vagus nerve for four minutes under anesthesia. Following stimulation, the animals were recovered for two hours, euthanized, and whole blood was collected through cardiac puncture. **B.)** Schematic of the charge-balanced stimulation waveforms used during stimulation, with short pulse width (top) and long pulse width (bottom). The actual waveform shapes used in this study are shown to the right.

While electrical stimulation is known to decrease serum TNFα in a setting of increased inflammation, its effect during normal physiology is unclear. To assess this, we delivered cervical stimulation to normal mice and measured serum cytokines two hours later. We found that specific combinations of stimulation parameters significantly changed levels of serum TNFα. Specifically, stimulation at the short pulse width (50 μs) at 30 Hz pulse and 200 μA amplitude produced a significant decrease in TNFα (Mann Whitney U=216, *P*<0.05; **Figure 2A**). Stimulation with a 50 μs pulse width at 100 Hz and 750 μA also resulted in a decrease in serum TNFα (Mann Whitney U=6, *P*<0.05; **Figure 2B**). When we increased the pulse width to 250 μs during stimulation, we observed a significant increase in serum TNFα levels at 30 Hz and 750 μA, compared to sham mice (Mann Whitney test, U=60, *P*<0.0001; **Figure 3A**). In contrast, stimulation 250 μs and 100 Hz produced no statistically significant changes in serum TNFα (Mann Whitney U=38, *P*=0.17; **Figure 3B**). These results suggest that specific stimulation parameters can alter serum TNFα in a bidirectional manner.

**Figure 2.**
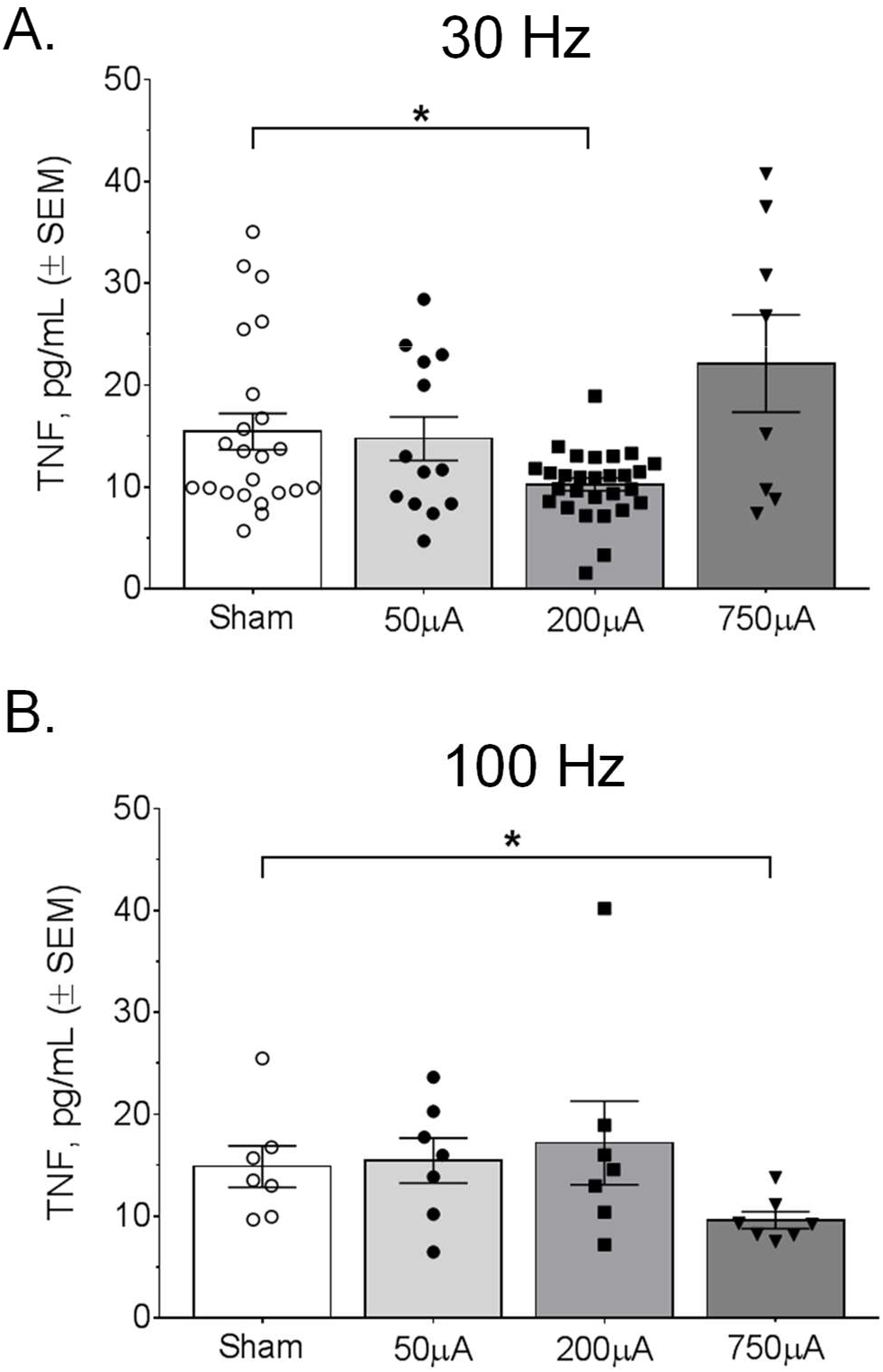
Specific stimulation amplitude and frequency combinations at 50 μs pulse widths reduce serum TNFα levels. **A.)** A significant decrease in TNFα, compared to the sham group, was observed with 30 Hz stimulation and a pulse amplitude of 200 μA. **B.)** A significant decrease in TNFα was observed at 100 Hz stimulation with a pulse amplitude of 750 μA.

**Figure 3.**
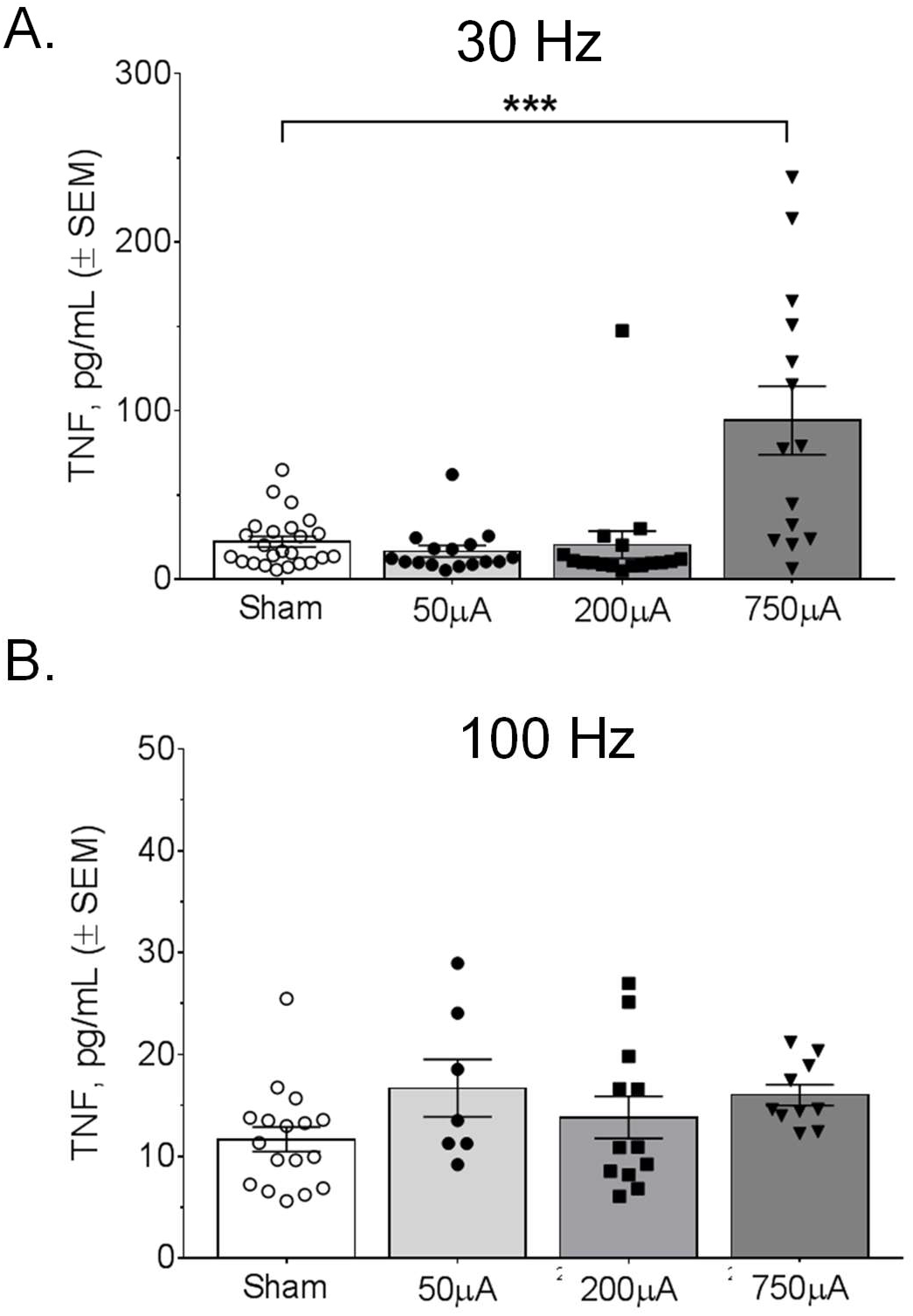
Serum TNFα is significantly increased by vagus nerve stimulation at 250 μs for a specific parameter combination. **A.**) Stimulation resulted in a significant increase in TNFα at 30 Hz and 750 μA pulse amplitude, compared to the sham group. **B.**) No significant changes in serum TNFα were observed with the 250 μs pulse width at 100 Hz.

IL-10 is a potent anti-inflammatory cytokine that suppresses Th1 cells, NK cells, and macrophages during infection (Couper et al., 2008; Hutchins et al., 2013). To examine whether electrical stimulation could also change levels of serum IL-10, we used specific combinations of frequency, pulse width, and amplitude followed by serum IL-10 measurements. Interestingly, the shorter 50 μs pulse width at 30 Hz produced statistically significant increases in serum IL-10 at both the 50 μA and 750 μA amplitudes (50 μA, Mann Whitney U=115, *P*<0.05; 750 μA, Mann Whitney U=53, *P*<0.05; **Figure 4A**). Meanwhile, a 50 μs pulse width at 100 Hz stimulation resulted in no significant changes in IL-10 levels (Mann Whitney U=34, *P*=0.96; **Figure 4B**). With the longer 250 μs pulse width, we observed statistically significant increases in IL-10 at 750 μA amplitude for both the 30 Hz (Mann Whitney U=53, *P*<0.05; **Figure 5A**) and 100 Hz frequencies (Mann Whitney U=18, *P*<0.001; **Figure 5B**). There was also a significant increase in IL-10 at 100 Hz with the 250 μs pulse width at 50 μA amplitude (Mann Whitney U=20, *P*<0.05; **Figure 5B**). This demonstrates that specific parameters have a different effect on serum IL-10, compared to TNFα.

**Figure 4.**
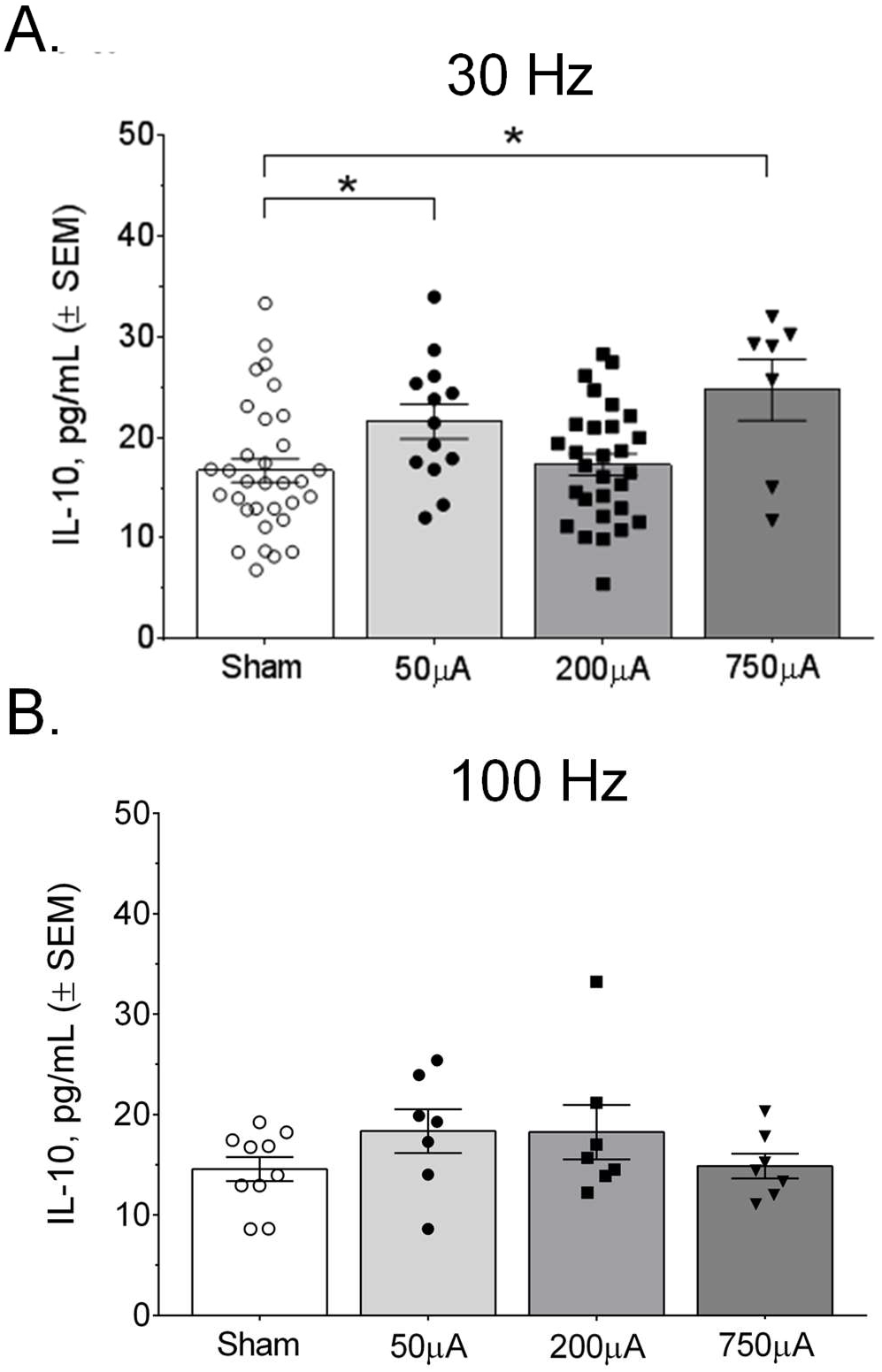
Serum IL-10 is increased by select parameters of electrical stimulation with 50 μs pulse width. **A.)** Stimulation for 50 μs at 30 Hz produced significant increases, compared to the sham group, in IL-10 for both 50 μA and 750 μA pulse amplitudes. **B.)** No changes in serum IL-10 were observed for 100 Hz stimulation across the four pulse amplitudes.

**Figure 5.**
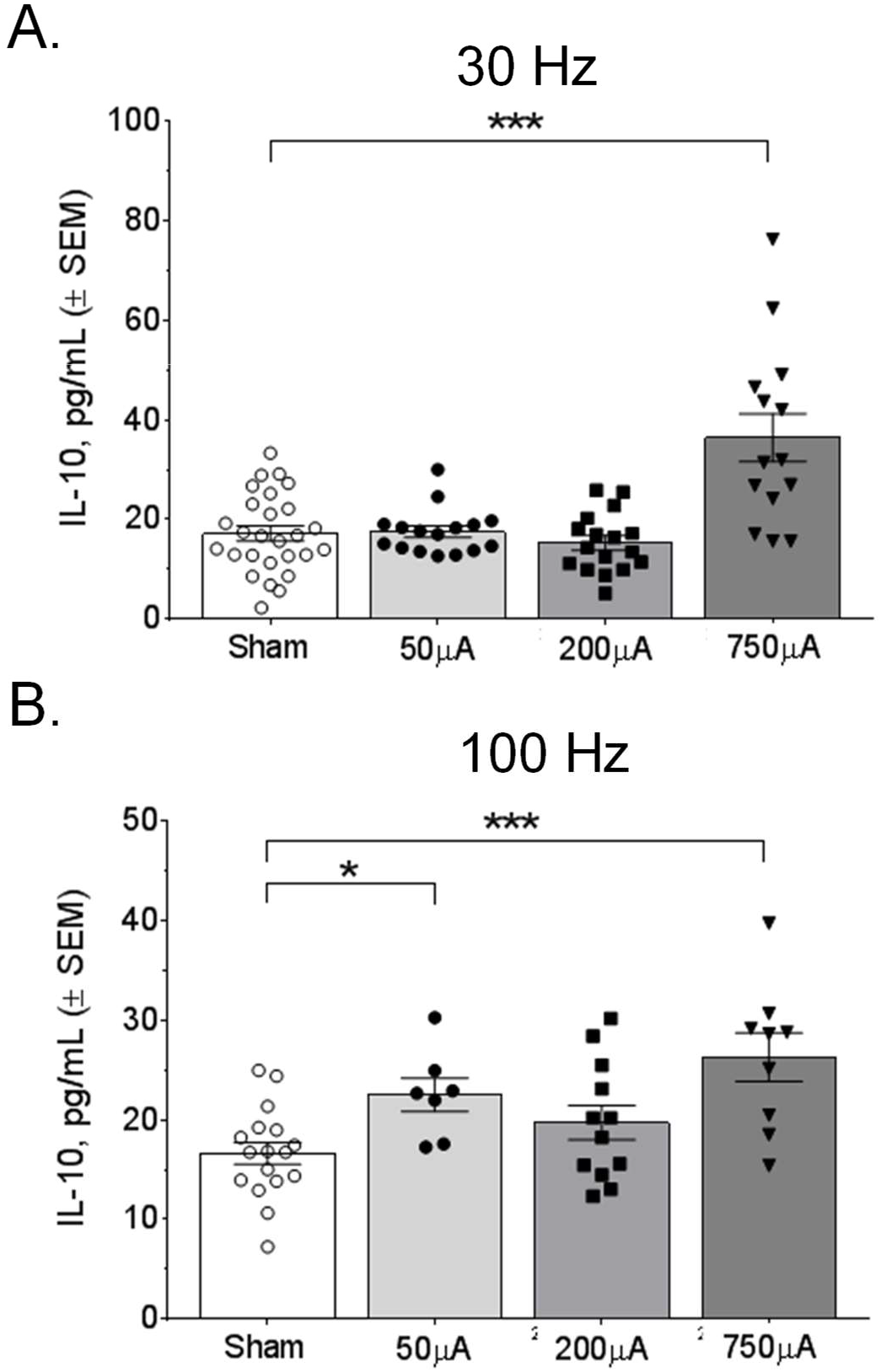
Nerve stimulation with 250 μs pulse width increased serum IL-10 at several different parameters. **A.)** Stimulation with 250 μs pulses at 30 Hz produced a marked increase in IL-10 at the 750 μA pulse amplitude. **B.)** Stimulation with 250 μs pulses at 100 Hz produced significant increases at both 50 μA and 750 μA pulse amplitudes.

Electrical nerve stimulation-induced bradycardia has been used to index nerve fiber activation during stimulation, specifically activation of the intermediate diameter B-fibers within the vagus nerve (Musselman et al., 2019; Yoo et al., 2013; Yoo et al., 2016). During our stimulation experiments, we measured heart rate with a continuous heart rate monitor to obtain an indirect measure of B-fiber recruitment. Stimulation with 50 μs pulses resulted in bradycardia only at the 750μA and 30Hz parameter (−17.1 ± 2.9% decrease; **Figure 6A**). The 50 μs pulse width stimulation did not produce bradycardia at 100 Hz for any of the tested pulse widths (**Figure 6B**). With the longer 250 μs pulse at 30 Hz, bradycardia was produced at both 200 μA and 750 μA pulse amplitudes (**Figure 6C**). At 100 Hz, stimulation with 250 μs pulse width produced the largest measured decrease in heart rate (−20.4 ± 4.4%; **Figure 6D**). These results indicate that alteration of stimulation parameters results in differential effects on heart rate, with longer pulse width increasing the likelihood of recruiting cardiac innervating B-fibers in the mouse vagus nerve.

**Figure 6.**
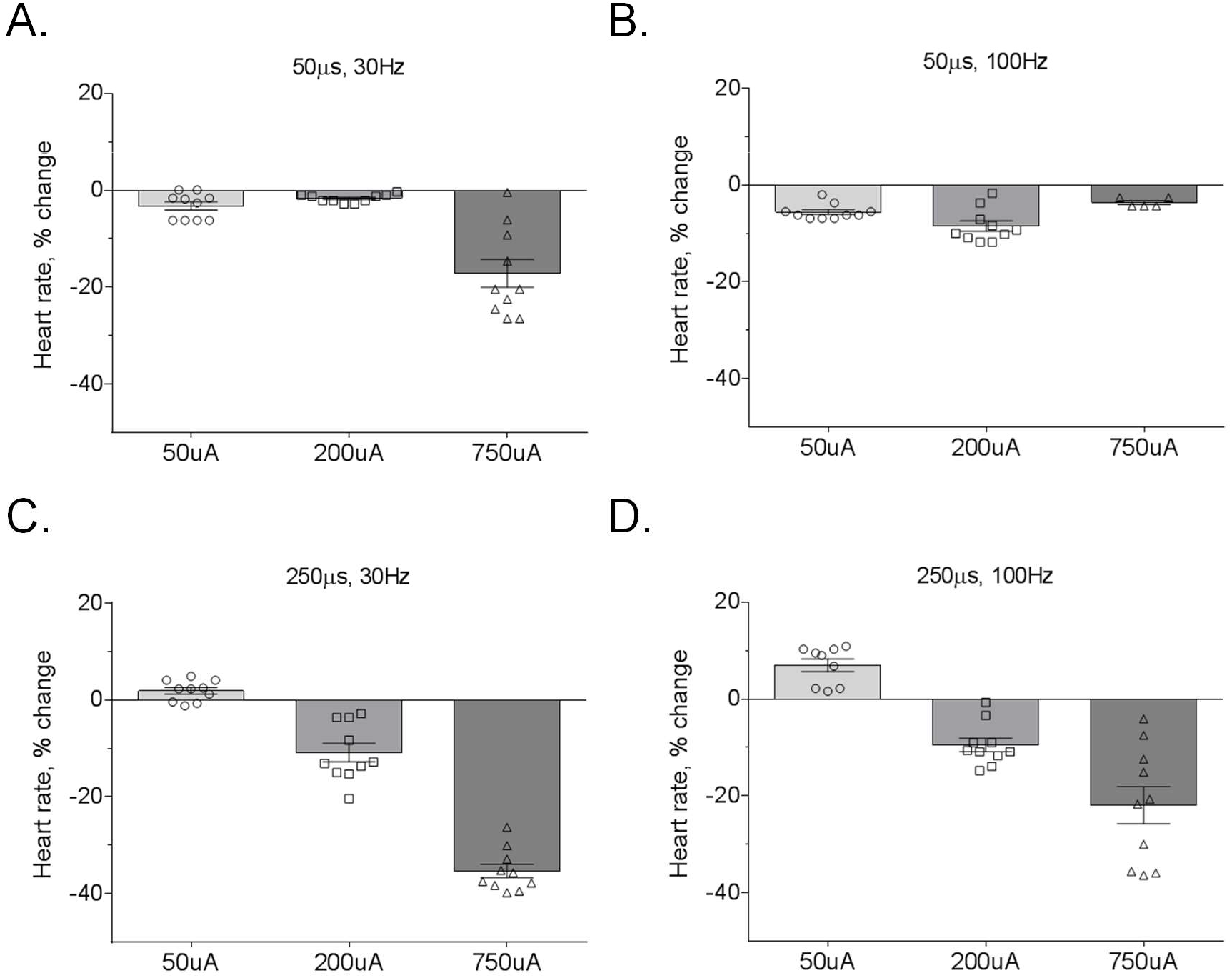
The effect of different vagus nerve stimulation parameters on heart rate. **A.)** Stimulation at 30 Hz with 50 μs pulse width resulted in bradycardia (≥10% reduction in heart rate) at only the 750 μA pulse amplitude. **B.)** At 100 Hz, bradycardia was not observed with the 50 μs pulse width. **C.)** Stimulation with the longer 250 μs pulse width resulted in bradycardia at 200 and 750 μA pulse amplitudes. **D.)** The largest decrease in heart rate was observed with the 250 μs pulse width at 100 Hz and 750 μA amplitude.

In addition to assessing changes in the prototypical pro-inflammatory cytokine TNFα and anti-inflammatory cytokine IL-10, we also measured the serum levels for a set of additional cytokines including: interferon gamma (IFN-γ), interleukin-12 p70 (IL-12 p70), interleukin-1 beta (IL-1β), interleukin-2 (IL-2), interleukin-4, (IL-4), interleukin-5 (IL-5), interleukin-6 (IL-6), and chemokine ligand C-X-C motif ligand 1 (CXCL1). We found that the serum levels of certain pro-inflammatory cytokines, such as IL-6, were increased across a wide range of stimulation parameters for both the short 50 μs and long 250 μs pulse widths (**Table 1, Table 2**). Serum CXCL1 was also significantly increased by stimulation across several parameter combinations. For anti-inflammatory cytokines, we did not observe changes in IL-4 for any stimulation parameter combination. These results suggest that certain cytokines are more susceptible to modulation via electrical stimulation than others. It should be noted that the measured concentrations of IFNγ, IL-1β, IL-2, IL-4, and IL-5 were very low across the sham and stimulation groups such that, while there may be statistically significant changes with some of the stimulation parameters, the low concentrations suggest a lack of physiological relevance for these specific cytokines (**Table 1, Table 2**).

**Table 1.** Cytokine levels (mean ± SEM) at 50μs pulse

| Cytokine | Sham | 50uA<br>30Hz | 200uA<br>30Hz | 750uA<br>30Hz | 50uA<br>100Hz | 200uA<br>100Hz | 750uA<br>100Hz |
| --- | --- | --- | --- | --- | --- | --- | --- |
| IFN-γ | 0.64 ± 0.15 | 0.48 ± 0.13 | 0.34 ± 0.05 | 0.46 ± 0.26 | 0.39 ± 0.11 | 0.53 ± 0.21 | 0.20 ± 0.02 |
| IL-12 p70 | 37.27 ± 5.72 | 34.18 ± 7.17 | 47.35 ± 6.77 | 30.35 ± 9.37 | 41.24 ± 11.16 | 58.15 ± 16.82 | 43.77 ± 10.46 |
| IL-1β | 1.79 ± 0.32 | 2.43 ± 0.41 * | 1.53 ± 0.21 | 2.03 ± 0.38 | 2.93 ± 0.54 | 2.15 ± 0.52 | 2.35 ± 0.73 |
| IL-2 | 1.75 ± 0.70 | 1.29 ± 0.65 | 1.82 ± 0.71 | 1.40 ± 0.35 | 2.25 ± 0.64 | 2.09 ± 0.88 | 0.36 ± 0.10 |
| IL-4 | 0.22 ± 0.08 | 0.37 ± 0.16 | 0.81 ± 0.41 | 0.47 ± 0.28 | 0.45 ± 0.18 | 0.43 ± 0.10 | 0.36 ± 0.20 |
| IL-5 | 3.12 ± 0.34 | 4.58 ± 0.89 | 4.02 ± 0.75 | 4.43 ± 0.88 | 5.38 ± 1.05 * | 4.42 ± 0.95 | 1.85 ± 0.20 * |
| IL-6 | 2289.58 ±<br>331.97 | 3270.73 ±<br>323.83 ** | 2653.11 ±<br>184.78 * | 5516.24 ±<br>1139.26 ** | 2826.10 ±<br>305.77 | 2921.93 ±<br>322.54 * | 2906.27 ±<br>320.22 * |
| CXCL1 | 739.68 ± 50.39 | 1004.54 ±<br>86.69 * | 1074.40 ±<br>60.44 *** | 786.88 ±<br>71.59 | 1131.30 ±<br>71.65 *** | 1025.21 ±<br>68.99 ** | 880.59 ±<br>92.48 |

Full multiplex panel results of serum cytokine levels following nerve stimulation with 50 μs pulse width. Values in red indicate significant differences from the Sham group. \*, $P < 0.05$ ; \*\*, $P < 0.01$ ; \*\*\* $P < 0.001$ .

**Table 2.** Cytokine levels (mean ± SEM) at 250μs pulse

| Cytokine | Sham | 50uA<br>30Hz | 200uA<br>30Hz | 750uA<br>30Hz | 50uA<br>100Hz | 200uA<br>100Hz | 750uA<br>100Hz |
| --- | --- | --- | --- | --- | --- | --- | --- |
| IFN-γ | 0.69 ± 0.16 | 0.58 ± 0.22 | 0.45 ± 0.09 | 0.96 ± 0.48 | 0.66 ± 0.20 | 0.51 ± 0.12 | 0.25 ± 0.03 |
| IL-12 p70 | 50.31 ± 15.11 | 48.26 ± 6.26 | 48.83 ± 13.70 | 39.63 ± 4.49 | 52.80 ± 13.92 | 41.90 ± 7.12 | 78.17 ± 9.33 ** |
| IL-1β | 3.10 ± 0.91 | 5.42 ± 2.04 ** | 2.01 ± 0.45 | 1.48 ± 0.34 | 4.07 ± 0.52 | 2.54 ± 0.44 | 2.15 ± 0.29 |
| IL-2 | 2.85 ± 1.12 | 1.69 ± 0.65 | 3.05 ± 1.23 | 1.88 ± 1.13 | 5.21 ± 2.31 | 4.80 ± 2.11 | 2.63 ± 0.56 |
| IL-4 | 0.34 ± 0.08 | 0.44 ± 0.22 | 0.38 ± 0.17 | 0.54 ± 0.29 | 0.67 ± 0.42 | 0.53 ± 0.21 | 0.20 ± 0.06 |
| IL-5 | 2.97 ± 0.36 | 4.36 ± 0.72 | 5.48 ± 1.26 * | 6.13 ± 1.41 * | 9.50 ± 2.74 ** | 6.19 ± 1.20 *** | 3.55 ± 0.65 |
| IL-6 | 2125.23 ±<br>243.76 | 2981.44 ±<br>376.14 * | 2911.30 ±<br>405.83 * | 5880.09 ±<br>1248.88 *** | 3541.48 ±<br>408.73 ** | 3040.72 ±<br>394.06 * | 5490.44 ±<br>473.28 **** |
| CXCL1 | 798.88 ±<br>55.16 | 947.72 ±<br>83.90 | 1134.26 ±<br>121.55 * | 907.32 ±<br>114.89 | 1192.14 ±<br>80.46 *** | 1717.80 ±<br>170.34 **** | 1081.02 ±<br>160.75 |

Full multiplex panel results of serum cytokine levels following nerve stimulation with 250 μs pulse width. Values in red indicate significant differences from the Sham group. \*, $P < 0.05$ ; \*\*, $P < 0.01$ ; \*\*\*, $P < 0.001$ ; \*\*\*\*, $P < 0.0001$ .

## DISCUSSION

The neural control of immunity through stimulation of the vagus nerve holds significant promise for treating inflammatory disorders. Here we have shown that a set of electrical stimulation parameters can change specific cytokine levels in the absence of inflammation. We found, to our surprise, that electrical stimulation of the vagus nerve with specific parameters in conditions of low circulating cytokines (i.e. healthy control mice) results in an elevation in serum levels of TNFα and IL-10, cytokines known to be involved in inflammatory responses. These results indicate that systemic cytokines respond to specific combinations of frequency, amplitude, and pulse width applied to the cervical vagus nerve. How these parameters affect the recruitment of specific fiber types within the vagus nerve remains a topic of important ongoing work, but our results indicate that the refinement of these parameters may be important in the neuromodulation of immunological responses mediated through vagal signaling.

The effect of vagus nerve stimulation on cytokines in the absence of systemic inflammation has not, to our knowledge, been carefully examined. This is the first demonstration that specific stimulation parameters can be used to increase serum TNFα and IL-10. TNFα is an inflammatory cytokine that is released rapidly by macrophages following infection and injury. Transient elevations in serum TNFα are required to coordinate host defense against pathogens and the repair of injured host tissue. While physiological inflammation is locally protective, pathological inflammation caused by persistent and unregulated TNF levels cause chronic tissue damage and drive the pathogenesis of inflammatory disease including rheumatoid arthritis, Crohn’s disease, and inflammatory bowel disease (Bonaz et al., 2016; Kalliolias 2016; Koopman et al., 2016). A large body of work established the critical role of TNFα in the pathogenesis of systemic inflammation in animal models and led to the clinical translation of TNFα blockade to treat chronic inflammation. The beneficial effect of vagus nerve stimulation for reducing TNFα in the context of inflammatory disorders is clear, however, an increase in proinflammatory cytokine levels may also be beneficial in certain physiological and pathologic contexts. An increase in circulating cytokines might be useful to bolster the immune system in conditions of immunosuppression, for example, due to immunodeficiencies or viral infection (Breen, 2002; Varzaneh et al., 2014). There is also evidence that certain cancer patients have lowered levels of certain circulating cytokines that might benefit from immunotherapy to increase those levels (Khan et al., 2018). As techniques are developed to better understand the relationship between vagus nerve activity and cytokine signaling (Steinberg et al., 2016; Tsaava et al., 2019; Zanos et al., 2018), there may also be other physiological conditions that would benefit from intentionally increasing cytokine levels.

Our results suggest that serum cytokines may likewise be controlled by specific fiber sets and firing patterns. Altering the parameters of the stimulation pulse affects the specific vagus nerve fibers that are recruited. A large body of evidence shows that increasing pulse width and amplitude, and consequently the charge delivered during the both the cathodic and anodic phases of the pulse, inhibits activation of large diameter fibers while selectively activating smaller diameter fibers (Baratta et al., 1989; Musselman et al., 2019). For example, lower amplitude stimulation activates A-fibers but as stimulation increases, the A-fiber activation becomes suppressed and B-fibers become activated to change heart rate (Burke et al., 1975). It is possible that the differential effect on heart rate may be a result of effective recruitment of large and small fiber types with the long pulse at lower amplitudes, and selective inhibition of larger fibers at the larger pulse widths. The effective pulse widths and stimulus amplitudes necessary to elicit heart rate changes is shown in **Figure 6**. As the stimulation charge increased, we generally observed more pronounced bradycardia, however, heart rate decreases were not always coupled to changes in serum cytokines. The largest increase in serum TNFα at 250 μs pulse width, 30 Hz, and 750 μA was associated with the largest decrease in heart rate, but TNFα levels did not change for two other stimulation parameters that induced bradycardia (**Figure 6**: 50 μs pulse width, 30 Hz, 750 μA and 250 μs pulse width, 100 Hz, 750 μA). This dissociation between bradycardia and serum cytokines levels confirms prior work indicating that cytokine effects are independent of the B-fiber activation that induces bradycardia (Huston et al., 2007).

We have considered the possibility that different durations of total stimulation may also be an important variable in cytokine-specific modulation via the vagus nerve. Further, while the applicability of these findings may be most relevant for disease conditions, these studies were carried out in healthy control animals. Therefore, they should be interpreted with caution if trying to extrapolate to specific disease conditions. The parameter space of electrical nerve stimulation is important because varying timing, duration, and amplitude should affect activation of different target organs and brain structures. Electrical stimulation delivered to cervical vagus nerve is broadly applied to all nerve fibers, including sensory afferents and motor efferents. It is thought that B-fibers mediate motor efferent control of the visceral organs. Because the vagus nerve innervates several major organs in the viscera, stimulation of this neve has the potential to treat a broad range of disorders ranging including but not limited to arthritis, myocardial infarctions, and metabolic disorders (Kong et al., 2012; Levine et al., 2014; Chang et al., 2019).

The unexpected results here indicate that it is possible to stimulate the production of TNFα and other cytokines using electrical stimulation of the cervical vagus nerve. As bioelectronic therapies continue to evolve for controlling inflammation, increasingly refined stimulation strategies will be needed to activate fibers of interest (Grill, 2015; Tsaava et al., 2019), while also minimizing the potential for the negative outcomes and off-target effects associated with vagus nerve stimulation. The effectiveness of these selective stimulation paradigms will depend on knowing which specific fiber populations mediate the neural regulation of inflammation and the optimal techniques for activating them. Our results show that varying stimulation parameters of frequency, amplitude, and pulse width holds promise for the specific regulation of cytokine responses in both health and disease.

